# Mapping Multiple Factors Mediated Chromatin Interactions to Assess Regulatory Network and Dysregulation of Lung Cancer-Related Genes

**DOI:** 10.1101/2022.03.15.481871

**Authors:** Yan Zhang, Jingwen Zhang, Wei Zhang, Mohan Wang, Shuangqi Wang, Yao Xu, Lun Zhao, Xingwang Li, Guoliang Li

**Author notes:** Correspondence: Guoliang Li. Equal contribution (Yan Zhang, Jingwen Zhang).

## Abstract

Studies on the lung cancer genome are indispensable for developing a cure for lung cancer. Whole-genome resequencing, genome-wide association studies, and transcriptome sequencing have greatly improved our understanding of the cancer genome. However, dysregulation of long-range chromatin interactions in lung cancer remains poorly described. To better understand the three-dimensional (3D) genomic interaction features of the lung cancer genome, we used the A549 cell line as a model system. The generated high-resolution data revealed chromatin interactions associated with RNA polymerase II (RNAPII), CCCTC-binding factor (CTCF), enhancer of zeste 2 polycomb repressive complex 2 subunit (EZH2), and histone 3 lysine 27 trimethylation (H3K27me3) using specific antibodies and long-read chromatin interaction analysis by paired-end tag sequencing (ChIA-PET). The EZH2/H3K27me3-mediated interactions further silenced target genes, either through loops or domains, and showed distributions along the genome distinct from and complementary to those associated with RNAPII. We found that cancer-related genes were highly enriched in chromatin interactions. We identified abnormal interactions associated with oncogenes and tumor suppressors, such as additional repressive interactions on *FOXO4* and promoter – promoter interactions between *NF1* and *RNF135*. Knockout of abnormal interactions reversed the dysregulation of cancer-related genes, suggesting that chromatin interactions are essential for proper expression of lung cancer-related genes. These findings demonstrate the 3D landscape and gene regulatory relationships of the lung cancer genome.

## Introduction

Lung cancer is currently one of the most lethal and common cancers [1]. A series of high-throughput sequencing studies have identified critical point mutations, structural variations, and copy number variations associated with tumorigenesis in lung cancer [2–5], greatly augmenting our understanding of the cancer genome. However, many risk-related single nucleotide polymorphisms (SNPs) and genomic variations are found in regulatory and intergenic regions [6]. Linking these distal regulatory elements to their target genes is crucial for a more comprehensive understanding of the cancer genome and the potential functions of noncoding regions.

Chromosome conformation capture (3C)-related technologies, such as chromatin interaction analysis by paired-end tag sequencing (ChIA-PET) [7], can be used to explore the three-dimensional (3D) genome structure and link distal regulatory elements to their target genes. Many mammalian diseases are associated with dysregulation of the spatial structure of the genome. For example, abnormal topologically associated domains (TADs) can lead to abnormal development and malformation by disrupting interactions among TADs [8]. Previous studies have used long-range interactions to map noncoding SNPs to target genes in patients with schizophrenia [9]. Dysregulation of cancer-related genes based on long-range interactions in cancer highlights the importance of spatial structures in maintaining normal cell function [10, 11]. Therefore, clarifying 3D genomic disorders in cancer cells may be an innovative way to conduct cancer research. A recent study used Hi-C to characterize the larger-scale 3D genomic structures of lung cancer [12]. However, 3D information related to gene expression regulation still needs to be more fully studied.

Polycomb repressive complex 2 (PRC2, which has core subunites EZH2, SUZ12, EED, and other components) is critical for normal development and silencing of remote genes [13–17]. A recent study on genome-wide interactions associated with the PRC2 complex in a mouse embryonic stem cell line suggested that the promoter – promoter interactions between developmentally regulated genes play an irreplaceable role in gene silencing [18]. Studies have also shown that histone 3 lysine 27 trimethylation (H3K27me3)-enriched domains have distinct intradomain interactions that play essential roles during development and in cancer cells [19, 20]. Thus, EZH2 and repressive histone modification H3K27me3 are potentially worthy factors for enriching genomic functional interactions.

In this study, we used A549 lung cancer cells and noncancerous BEAS-2B epithelial cells as model systems and applied the long-read ChIA-PET [21] approach to map global interactions associated with different factors, such as RNA polymerase II (RNAPII), CCCTC-binding factor (CTCF), EZH2, and H3K27me3. By studying the patterns and functions of different categories of interactions and comparing the differences in interactions between lung cancer and noncancerous cells, we discovered the mechanisms of dysregulation at the 3D genome level, providing new insights into the transcriptional regulation and genomic characteristics of human lung cancer.

## Results

### Multiple factors mediated chromatin interactions in A549 lung cancer cell line

To investigate the 3D genome architecture and its regulatory function in lung cancer, we performed long-read ChIA-PET associated with CTCF, RNAPII, EZH2, and H3K27me3 in the lung cancer cell line A549 and noncancerous cell line BEAS-2B. These factors were selected because CTCF represents a classic structural protein in the 3D genome, RNAPII is associated with active gene transcription, and EZH2 and H3K27me3 are associated with repressive chromatin [18, 20, 22, 23]. These different chromatin interactomes should help clarify the 3D genome of the A549 lung cancer cell line. We obtained ∼375 million uniquely mapped paired-end reads and ∼283 000 chromatin loops from six ChIA-PET libraries (Table S1). The biological replicates for each factor-mediated ChIA-PET showed excellent reproducibility. The correlations among contact matrices and visible clusters are shown in Figure S1. We combined interaction pairs of the four factors and constructed a contact heatmap at the chromosomal level. The 1-Mb and 100-kb resolution contact heatmaps of chromosome 14 showed chromosome interaction patterns, where the A/B compartments and TADs can be identified (**Figure 1A and B**). We ascertained the distinct features of the interactions associated with each factor (Figure 1C), RNAPII-mediated chromatin interaction loops were mainly located in active regions, while EZH2- and H3K27me3-mediated chromatin interaction loops were mainly located in repressive regions. Moreover, the EZH2- and H3K27me3-associated interaction loops were mainly seen between their high peaks and broad repressed regions, showing marked differences from the interaction loops of RNAPII.

**Figure 1.**
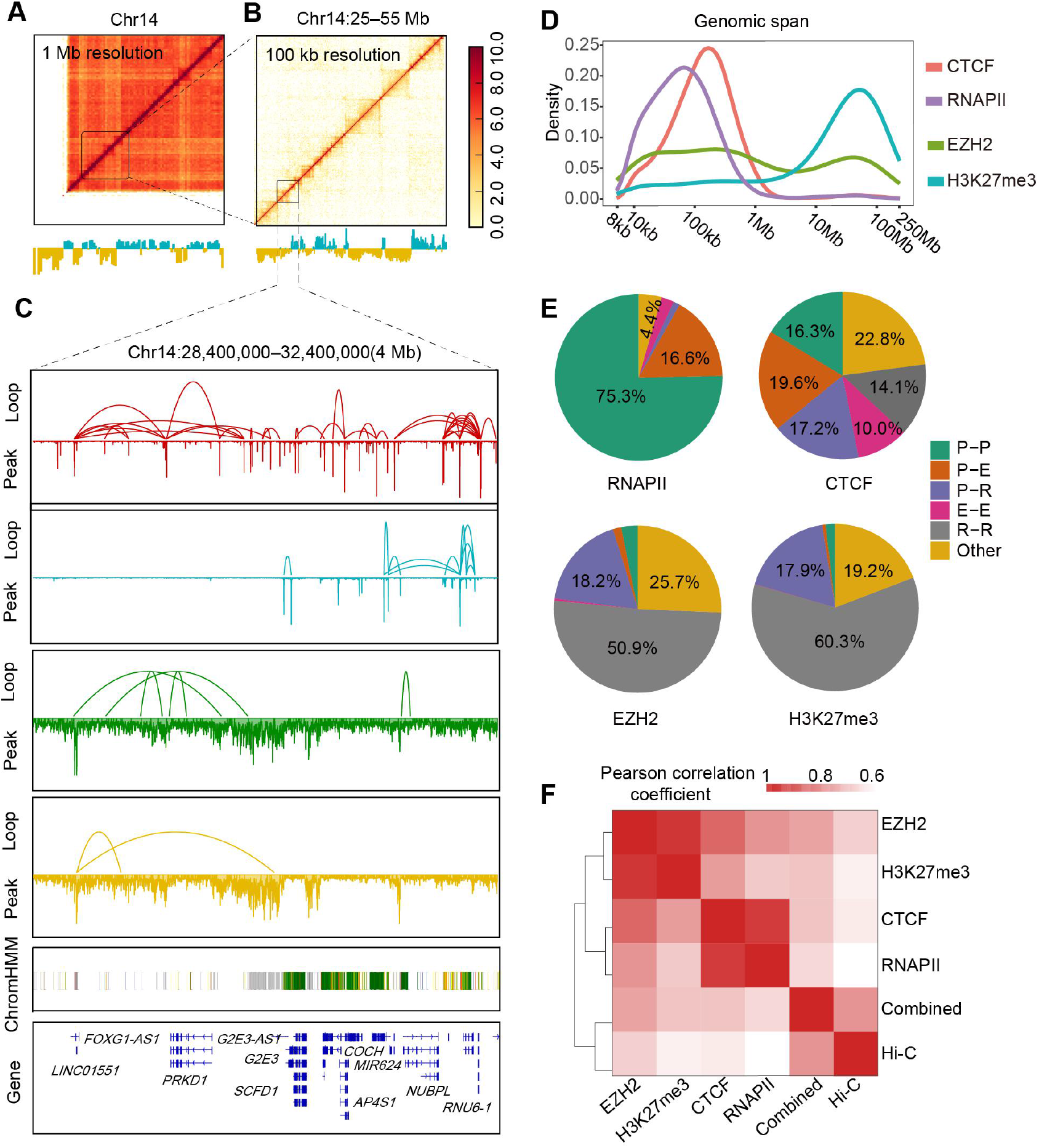
ChIA-PET interactions mediated by four different factors in A549 lung cancer cell line. **A-B.** Chromatin interaction heatmaps and A/B compartments of chromosome 14 in A549 cell line. Combined ChIA-PET interaction heatmaps based on CTCF, RNAPII, EZH2, and H3K27me3 ChIA-PET data at 1-Mb resolution (**A**) and 100-kb resolution (**B**). Lower panels show A (blue) and B (yellow) compartments calculated from principal component analysis. **C.** Loop and peak views of ChIA-PET data in 4-Mb region on chromosome 14 at zoom-in scope. For each data track, loop view is at the top, and peak view is at the bottom. Chromatin state annotation by ChromHMM was obtained from the Roadmap Epigenomics Mapping Consortium [52]. **D.** Different genomic spans of loops in ChIA-PET associated with CTCF, RNAPII, EZH2, and H3K27me3. **E.** Pie charts of different interaction categories (promoter (P)-promoter, promoter – enhancer (E), promoter – repressor (R), and others) from ChIA-PET mediated by four different factors. **F.** Heatmap of Pearson correlation coefficients between individual-factor mediated ChIA-PET interactions for CTCF, RNAPII, EZH2, H3K27me3, combined data, and Hi-C data.

We plotted the span distributions of the loops mediated by different factors (Figure 1D). In general, the RNAPII loops showed the shortest span and the highest density of loop spans was less than 100-kb. The CTCF loops showed the highest span density slightly over 100-kb. While the span distribution of EZH2 loops showed two peaks, the shorter one was less than 1-Mb, and the longer one was between 10 Mb and 100 Mb. And H3K27me3 loop span showed the hightest density between 10 Mb and 100 Mb. Jodkowska and colleagues observed a similar distribution of chromatin interactions with promoter-capture Hi-C, although the associated proteins were not reported in their study [24]. Subsequently, we described the distribution of interactomes associated with different factors on gene regulatory elements. As shown in Figure 1E, significant differences were found in pairs of regulatory elements between these factors, especially in terms of the proportion of promoter-involved interactions. Of the RNAPII-mediated loops, 92% were promoter – promoter or promoter–enhancer loops. In comparison, only a small proportion (5% for EZH2 and 2% for H3K27me3) of active promoter-centered interactions was organized by EZH2 and H3K27me3. For the two repressive factors, more than 70% of the interactions were repressor-associated interactions. Moreover, the loops associated with CTCF were mostly evenly distributed in the different categories, confirming the viewpoint that CTCF is more involved in structural maintenance than in gene regulation. The Pearson correlation coefficient (PCC) matrices of the four interactomes and Hi-C are shown in Figure 1F. The interaction matrices of EZH2 and H3K27me3 showed high similarity, CTCF and RNAPII showed high similarity, and the combined interaction matrices of the four factors were highly similar to Hi-C, suggesting that the combined interactomes of these factors reflected the whole-genome interactions observed with Hi-C. Therefore, comprehensive analysis of different factors associated chromatin interactions with distinct features can provide details on the 3D genome architecture and clarify the regulatory relationship of the A549 lung cancer cell line.

### High-resolution chromatin interaction analysis can reveal hierarchical genomic structures

Chromatin fragmentation by sonication and factor enrichment by antibodies in ChIA-PET can increase the resolution of interactive maps and the accuracy of functional elements. Based on ChIA-PET data, we observed hierarchical genomic structures at different scales. In addition to the A/B compartments and TAD-like structures (Figure 1A and B), we obtained much finer structures from the ChIA-PET data. By examining the span of the paired-end reads, we observed that DNA distance distribution was periodic. For example, we detected 10-bp (**Figure 2A**) and 190-bp periods (Figure 2B), reflecting the turn of the DNA double helix and the single nucleosome structure, respectively. The interaction positions were not randomly distributed around the nucleosomes. Moreover, 400–700-bp periods were observed in the chromatin transitional regions (CTRs, regions between active and inactive regions, defined as the center 2 kb of the overlapping regions of the H3K9me3 and H3K4me3 ChIP-seq peaks) in the ChIA-PET data associated with H3K27me3 but were not observed in the non-CTRs (Figure 2C and 2D). The structure with 3–4 nucleosomes in the CTRs may be unique, as described previously [25], and may be a basic structure in chromosome fiber assembly. Therefore, our high-resolution ChIA-PET data showed hierarchical 3D genome structures at different scales (Figure 2E).

**Figure 2.**
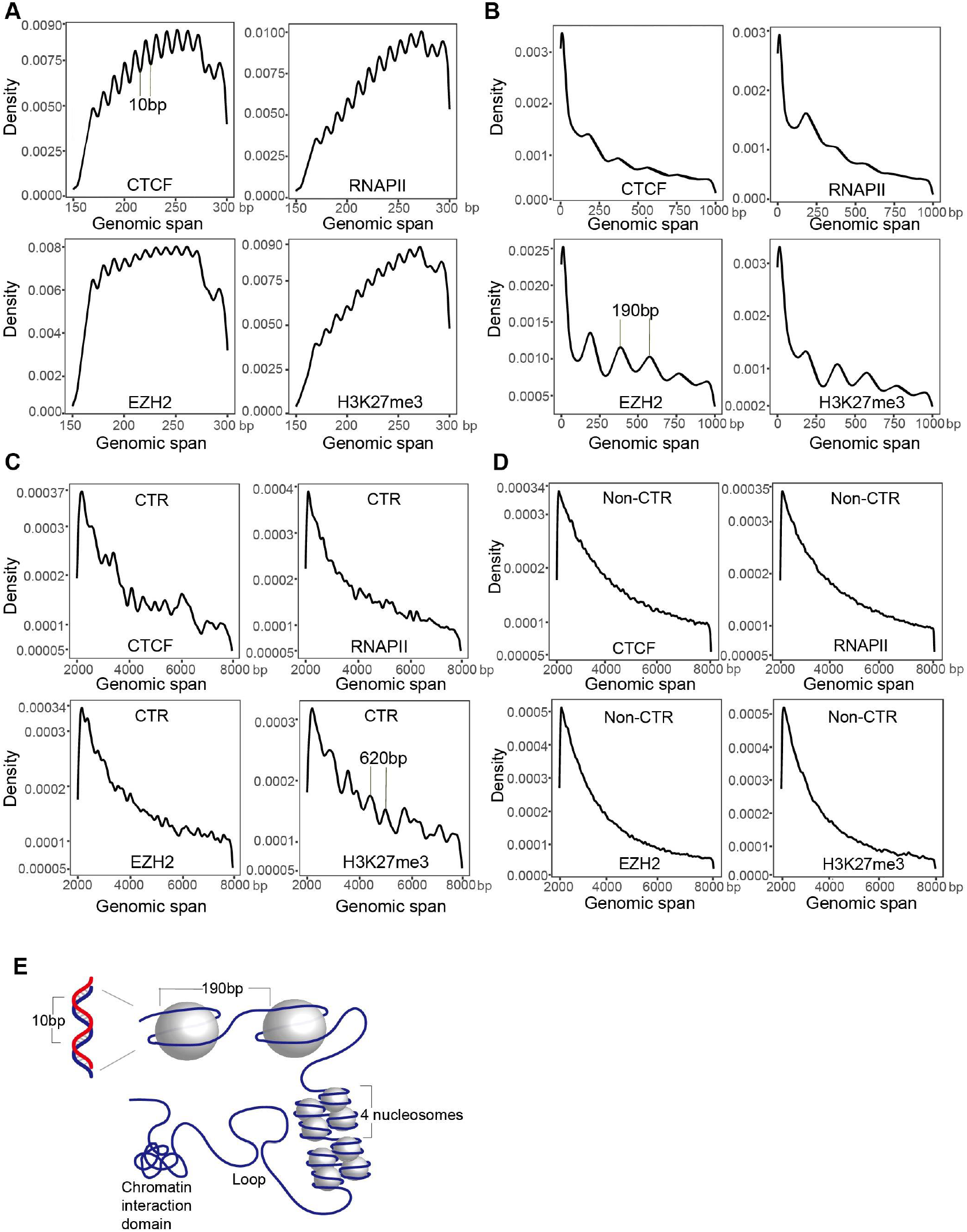
Hierarchical 3D genome structures obtained by ChIA-PET data. **A.** Distribution of paired-end tags (PETs) spanning 150 bp to 300 bp in ChIA-PET self-ligation data. X-axis represents PET spans in bp. The 10-bp period may represent DNA double helix. **B.** Distribution of PETs spanning 0 bp to 1 kb. The 190-bp span may represent mononucleosome structure. **C.** Distribution of PETs spanning 2 kb to 8 kb. Spans of 400–700 bp representing tetranucleosomes can be found in chromatin transitional regions (CTRs), especially in H3K27me3 ChIA-PET data. We defined CTRs as regions with overlapping H3K9me3 and H3K4me3 ChIP-seq peaks. There were no prominent periodic peaks in density curves in non-CTRs. **D.** Model and example of loop and loop domain structures revealed by A549 ChIA-PET data.

### EZH2- and H3K27me3-mediated chromatin interactions present distinct patterns and transcriptional activities from RNAPII

Interactions mediated by RNAPII and CTCF have been described in several studies [22, 23], and RNAPII-mediated promoter– promoter interactions can link gene pairs with strong coexpression patterns [23]. However, the characteristics and profiles of repressive interactions in lung cancer remain to be studied. Here, we determined whether genes with EZH2 or H3K27me3 promoter – promoter interactions are also transcriptionally coordinated. RNA sequencing (RNA-seq) data showed that most of the paired genes with RNAPII promoter–promoter interactions were highly expressed. Paired genes showing CTCF promoter – promoter interactions displayed weaker coexpression tendencies. In contrast, the expression levels of paired genes showing EZH2 or H3K27me3 promoter – promoter interactions were highly variable and exhibited much weaker coexpression relationships (**Figure 3A**). Low-expression pairs were much more abundant in repressive interactions than in RNAPII interactions. To further assess the coordinated transcription of paired genes across different conditions, we performed Pearson correlation analysis using the Cancer Genome Atlas (TCGA) lung adenocarcinoma (LUAD) RNA-seq data. Results indicated that genes involved in RNAPII promoter – promoter interactions were highly correlated, while those involved in promoter– promoter interactions with other factors were lowly correlated (Figure S2A). These analyses indicate that most gene pairs involved in RNAPII promoter – promoter interactions tend to be highly cooperatively transcribed. In contrast, gene pairs involved in EZH2 or H3K27me3 promoter–promoter interactions tend to be weakly (or not at all) cooperatively transcribed. In previous ChIP-Seq or GWAS SNP analyses, regulatory elements are assumed to regulate their nearest genes [26, 27]. However, this may not always be true. We observed that more than 64% of enhancers and 87% of repressors did not interact with their nearest promoters, instead bypassing several intervening genes to reach their target promoters (Figure S2B). In addition, proximal interactions, whether active or repressive, had stronger regulatory effects on target genes than distant interactions (Figure S2C).

**Figure 3.**
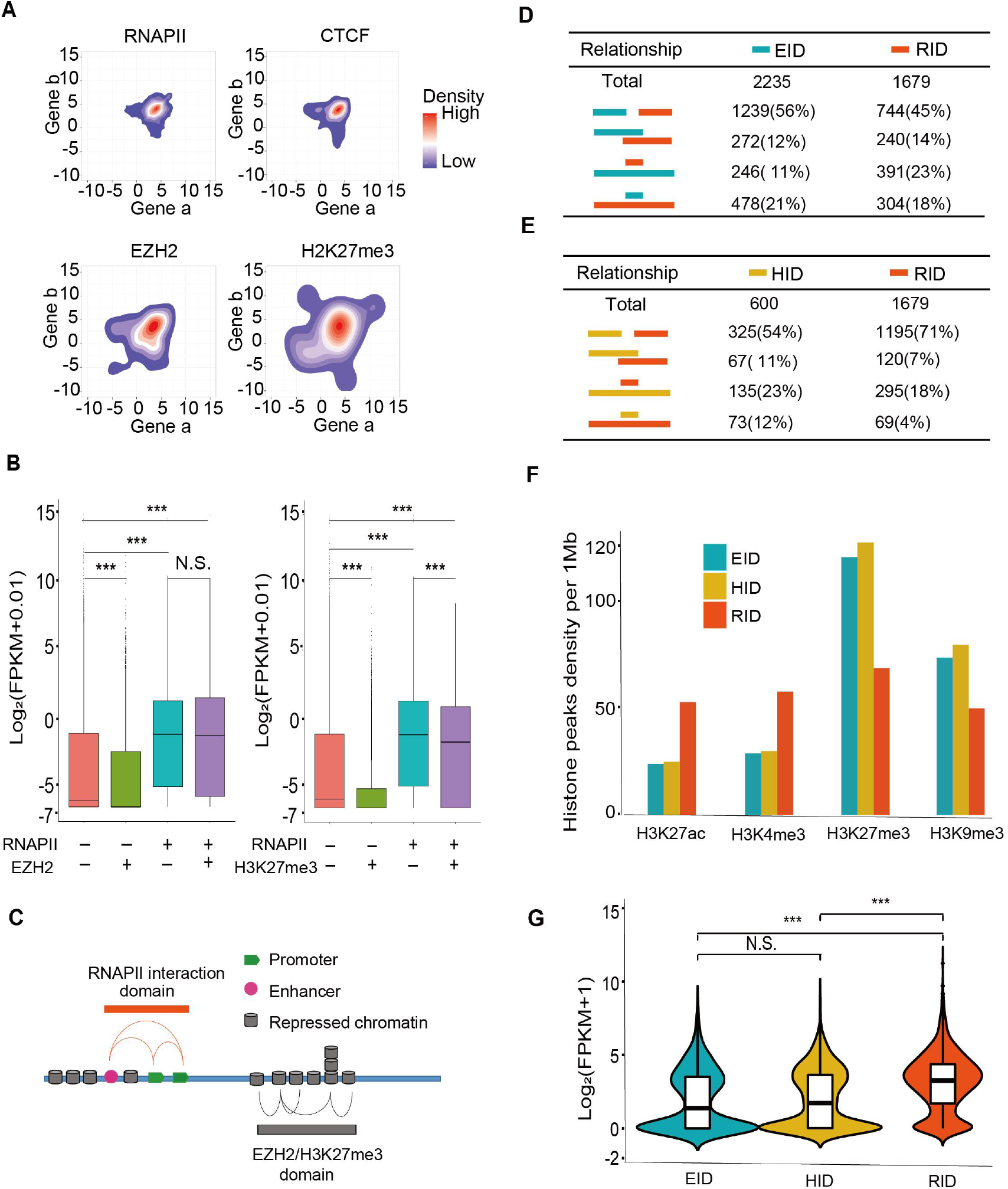
Characterization of four factors mediated chromatin interactions. **A.** Contour plot of log2-transformed RNA-seq FPKM values for promoter–promoter interacting genes in A549 cells. Gene pairs with RNAPII and CTCF interactions show concentrated high gene expression levels, while gene pairs with EZH2 and H3K27me3 interactions show more dispersed gene expression levels. **B.** Boxplots show gene expression with or without interactions mediated by different factors. *P*-value was determined using one-sided Mann-Whitney U test. *** *P* < 0.001; N.S., no significant difference. “+”, genes have loops mediated by this factor on gene promoter (−1 kb to +1 kb of transcription start site). “−”, no loops of this factor on gene promoter. **C.** Model of interaction domain is defined. Relatively high proportion and continuous loops for a factor form domain enriched in specific factor interactions. RID, RNAPII-associated interaction domain; EID, EZH2-associated interaction domain; HID, H3K27me3 (heterochromatin)-associated interaction domain. **D.** Table of relative positions and proportions of EIDs and RIDs. Proportions of independent, partially intersecting, and inclusive relationships of different interaction domains are shown in the table. **E.** Table of relative positions and proportions of HIDs and RIDs. **F.** Densities of histone modification peaks per 1 Mb in EIDs, HIDs, and RIDs. **G.** Violin plots overlaid with boxplots of gene expression levels in three kinds of interaction domains. *P-*value was determined using one-sided Mann-Whitney U test. *** *P* < 0.001; N.S., no significant difference.

We further investigated the effects of RNAPII and EZH2 loops on gene transcription. Results indicated that genes in the EZH2 loop anchors, but not in the RNAPII loop anchors, had the lowest expression levels, with levels lower than those of genes in neither loop anchors. In contrast, genes in the RNAPII loop anchors showed high expression levels. A similar trend was found when comparing the expression levels of genes with RNAPII and H3K27me3 interactions (Figure 3B). We also found that promoters bound to EZH2/H3K27me3 loop anchors with peaks were more repressed than promoters only bound to peaks, only bound to loop anchors, or neither (Figure S2D). These results suggest that EZH2- and H3K27me3-mediated loops can further silence target genes.

To characterize the spatial relationships between RNAPII-, EZH2-, and H3K27me3-associated chromatin topologies, we defined chromatin interaction domains based on loop connectivity and contact frequency (i.e., RID: RNAPII interaction domain; EID: EZH2 interaction domain; HID: H3K27me3 interaction domain) (Figure 3C, Figure S2E). In total, we identified 1 679 RIDs, 2 235 EIDs, and 600 HIDs. Most RIDs (71% and 45%, respectively) were isolated from HIDs and EIDs, while 23% of RIDs were contained in EIDs and 18% of RIDs were contained in HIDs (Figure 3D and E). We also investigated the epigenomic marks and transcriptional activity of different chromatin interaction domains. We found that EIDs and HIDs exhibited lower densities of active histone marks (H3K27ac and H3K4me3) and higher densities of inactive histone marks (H3K27me3 and H3K9me3). In contrast, RIDs presented the opposite histone properties (Figure 3F). In addition, EIDs and HIDs contained fewer active genes (Figure S2F) and exhibited lower transcriptional activities (Figure 3G) than RIDs. These findings suggest that the interactions associated with different factors exhibit distinct distributions and coexpression features in the genome, and that factor-specific chromatin interaction domains represent distinct epigenomic properties and transcriptional activities.

### High-resolution loops map whole-genome regulatory relationship of cancer-related genes and SNPs

Genomic interactions can reveal the spatial proximity and regulatory events of genomic sites [23, 28, 29]. Interaction mapping of cancer-related genes may increase our understanding of the regulatory elements of these genes. ChIA-PET loops can directly link distal elements to the promoters of target genes to yield regulatory information and can accurately identify specific transcription start sites (TSSs). Here, we identified genes and noncoding regions interacting with lung cancer-related genes (including 203 oncogenes and tumor suppressor genes from the Network of Cancer Genes (NCG) [30] and 100 survival-related genes from the Gene Expression Profiling Interactive Analysis (GEPIA) database [31]) using high-resolution interaction data (Table S2). Several typical cancer-related genes exhibited interactions mediated by CTCF, EZH2, H3K27me3, and especially RNAPII (**Figure 4A**, Table S2), thus suggesting a regulatory relationship with these genes. As seen in Figure 4B, the oncogene *ERBB2* showed several strong RNAPII interactions with other genes or noncoding regions. Compared with neighboring or distant genes not in the interaction domain, genes showing *ERBB2* interactions demonstrated obvious coexpression patterns (Figure 4C), though similar coexpression patterns were not observed in normal samples (Figure S3A). For *ERBB2*, interaction site mutation or dysregulation may be a new target for cancer studies. The coexpression pattern between *ERBB2* and other genes, such as *PGAP3*, *STARD3*, and *PNMT*, may be a marker of cancer. Another example of interaction mapping is the survival-related gene *DKK1*, which was not involved in EZH2 interactions and showed higher expression in the A549 cell line than in BEAS-2B cell line (Figure S3B).

**Figure 4.**
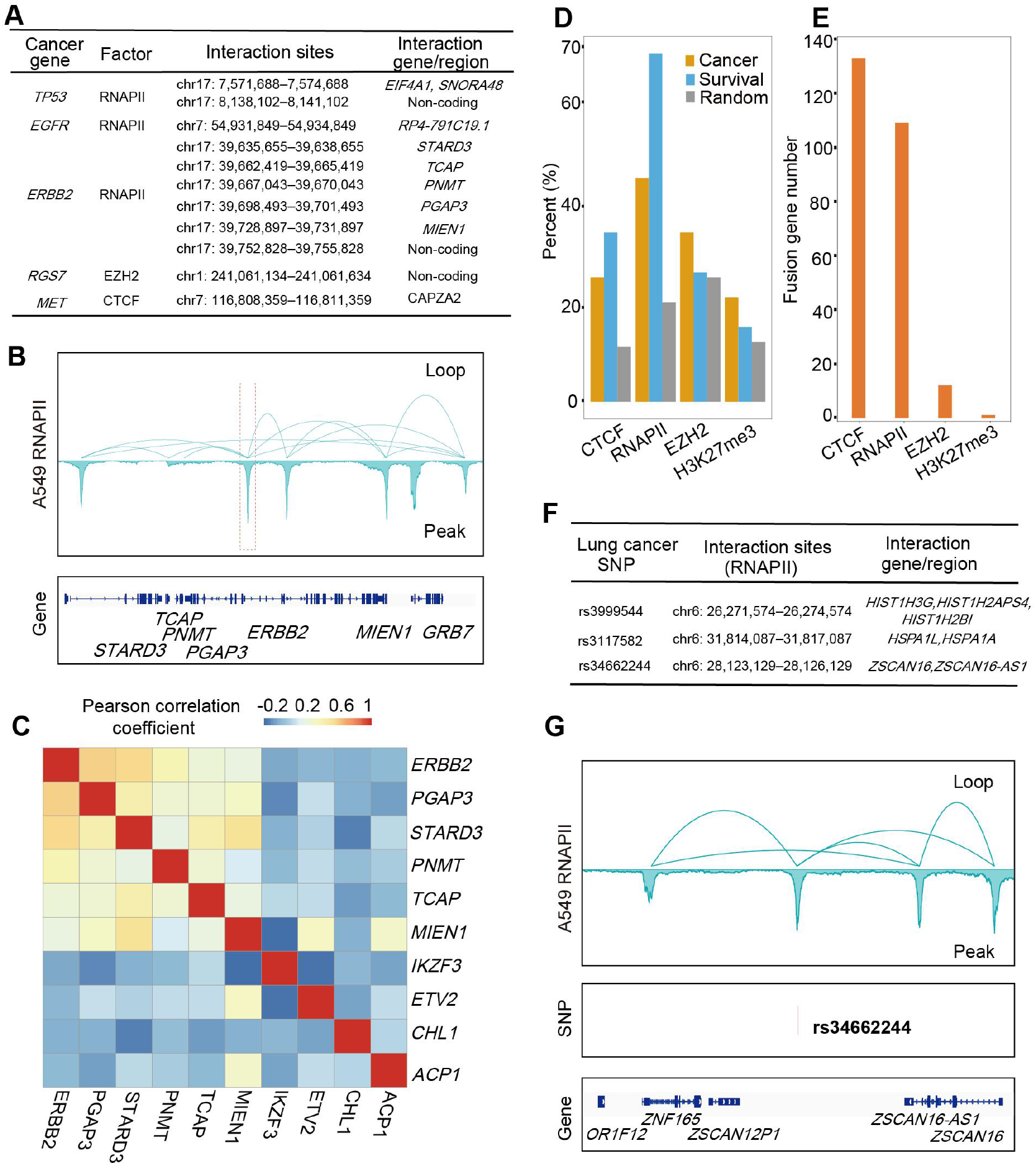
ChIA-PET interaction pairs show regulatory relationship of cancer risk-related genes and SNPs. **A.** Typical regions or genes interacting with cancer-related genes. If a gene promoter was located in the interacting region, it was defined as an interacting gene. Noncoding means that the interacting region is an intergenic region far away from gene body. **B.** enhancer – promoter or promoter – promoter interactions around oncogene *ERBB2* promoter. **C.** Pearson correlation coefficients between genes in TCGA LUAD tumor samples. *ERBB2* is a lung cancer-related gene. *STARD3*, *TCAP*, *PNMT*, *PGAP3*, and *MIEN1* are linked with ERBB2 by RNAPII loops. They are highly coexpressed with *ERBB2*. *IKZF3* is a gene near *ERBB2* and is not linked with *ERBB2*. *ETV2* is an inter-chromosome linking gene related to *ERBB2*. *CHL1* and *ACP1*, two randomly selected genes located on other chromosomes without interactions with *ERBB2*, were used as negative controls. **D.** Proportions of cancer- and survival-related genes involved in remote interactions mediated by four factors. “Random” represents 100 genes randomly selected in genome. **E.** Number of fusion transcripts overlapping with interaction loops associated with different factors. In total, 854 gene pairs from TCGA fusion transcripts and 854 random gene pairs were analyzed. Random gene pairs did not overlap with any loops. **F.** Selected regions or genes interacting with risk SNPs in lung cancer. **G.** Example of enhancer – promoter chromatin interactions linking one SNP rs34662244 in an intergenic region in chromosome 6 and its target genes. Upper track shows interaction clusters associated with RNAPII around SNP rs34662244. According to ChIA-PET data, *ZSCAN 16*, *ZSCAN 16-AS1*, and *ZNF165* are target genes of this SNP.

Interestingly, lung cancer-related genes were highly enriched in the RNAPII and CTCF interaction anchors (Figure 4D). Notably, nearly half were found to participate in chromatin interactions, with greater enrichment than randomly chosen genes. This enrichment suggests that mapping the interactomes of these genes would aid in studying the potential carcinogenic function of these cancer-related interactions in lung cancer. In addition, the anchor region may be more likely than other regions to significantly affect the genome, thus resulting in a cancer phenotype.

In lung cancer samples, tumor-specific fusion transcripts are suggested to disrupt normal gene function or activate proto-oncogenes, thus driving tumorigenesis [32–34]. We assessed the TumorFusions database list [35] in TCGA LUAD samples to determine whether long-range interactions exist in the spatial genome between gene pairs. Among the 854 fusion host gene pairs, more than 100 had loops mediated by RNAPII or CTCF (Figure 4E). In contrast, only a few pairs had interactions mediated by EZH2 or H3K27me3. As a control, 854 random pairs of expressed genes showed very low probabilities of participating in the interactions. These results suggest a positive relationship between TCGA fusion transcripts and spatial contacts, in which interactive loops may be potential facilitators of fusion transcripts.

We also mapped the interaction sites of high-risk SNPs in lung cancer (Figure 4F, Table S2). The lung cancer risk SNP rs34662244 was located in the anchor of two enhancer – promoter interaction clusters with target genes *ZNF165*, *ZSCAN16*, and *ZSCAN16-AS1* (Figure 4G). These interaction clusters suggest a potential role of this noncoding SNP in regulating target genes over a long distance. Moreover, based on all-gene expression comparison of TCGA LUAD tumor samples and normal samples, we found that a large proportion of SNP-interacting genes had higher expression levels in lung cancer samples than in normal samples (Figure S3C), suggesting a potential cancer driver function of SNP-associated genes. Therefore, our high-resolution chromatin interaction landscape can provide regulatory information of lung cancer-related genes and SNPs.

### Abnormal chromatin interactions are associated with dysregulation of oncogenes and tumor suppressor genes

As RNAPII- and EZH2-associated interactions showed significant regulation of gene expression, we analyzed whether abnormal expression of oncogenes or tumor suppressor genes is related to abnormal interaction patterns in lung cancer. We conducted joint analysis of differential gene expression and differential interactions between the lung cancer cell line A549 and noncancerous cell line BEAS-2B (Figure S4A and B). For genes associated with RNAPII chromatin interactions, the A549 and BEAS-2B cells showed considerable similarity. The differentially expressed and A549-specific anchor genes were significantly enriched in the Gene Ontology (GO) biological process term “anterior/posterior pattern specification”. The genes enriched in this term were mainly lung cancer-related *HOXB* gene clusters on chromosome 17 (Figure S4C). In the A549 cells, tandem gene clusters formed an RNAPII interaction domain and had higher binding peaks and gene expression than in BEAS-2B cells. The differentially expressed and BEAS-2B-specific anchor genes were significantly enriched in “DNA binding” (Figure S4D), and most belonged to the *ZNF* gene family and functioned as transcription factors. Thus, the *ZNF* family may play a role in lung cancer tumorigenesis.

Among abnormal interaction-associated genes, *RNF135* and *NF1* are two adjacent genes located on chromosome 17 [36]. The *NF1* gene is a repressor of the RAS signaling pathway, and mutation or abnormal expression of *NF1* may lead to lung cancer [37, 38]. Studies have also suggested that *RNF135* may exhibit oncogenic functions in glioblastoma [39]. Here, based on RNA-seq differential expression analysis, *NF1* and *RNF135* showed significantly higher expression in the A549 cells than in the BEAS-2B cells (fragments per kilobase per million mapped reads [FPKM] of *NF1* in A549: 35.3, BEAS-2B: 13.2; FPKM of *RNF135* in A549: 11.9, BEAS-2B: 6.0) (Table S3). In the A549 cells there was a strong RNAPII mediated promoter – promoter interaction cluster between these two genes (**Figure 5A**), which was not observed in the BEAS-2B cells. promoter–promoter interaction seems to increase the expression of this gene pair. In TCGA database analysis, both *NF1* and *RNF135* showed higher expression levels and larger variance in the LUAD tumor samples than in the normal samples (Figure 5B). In the survival analysis, *RNF135* overexpression indicated poor outcome in patients (Figure 5C).

**Figure 5.**
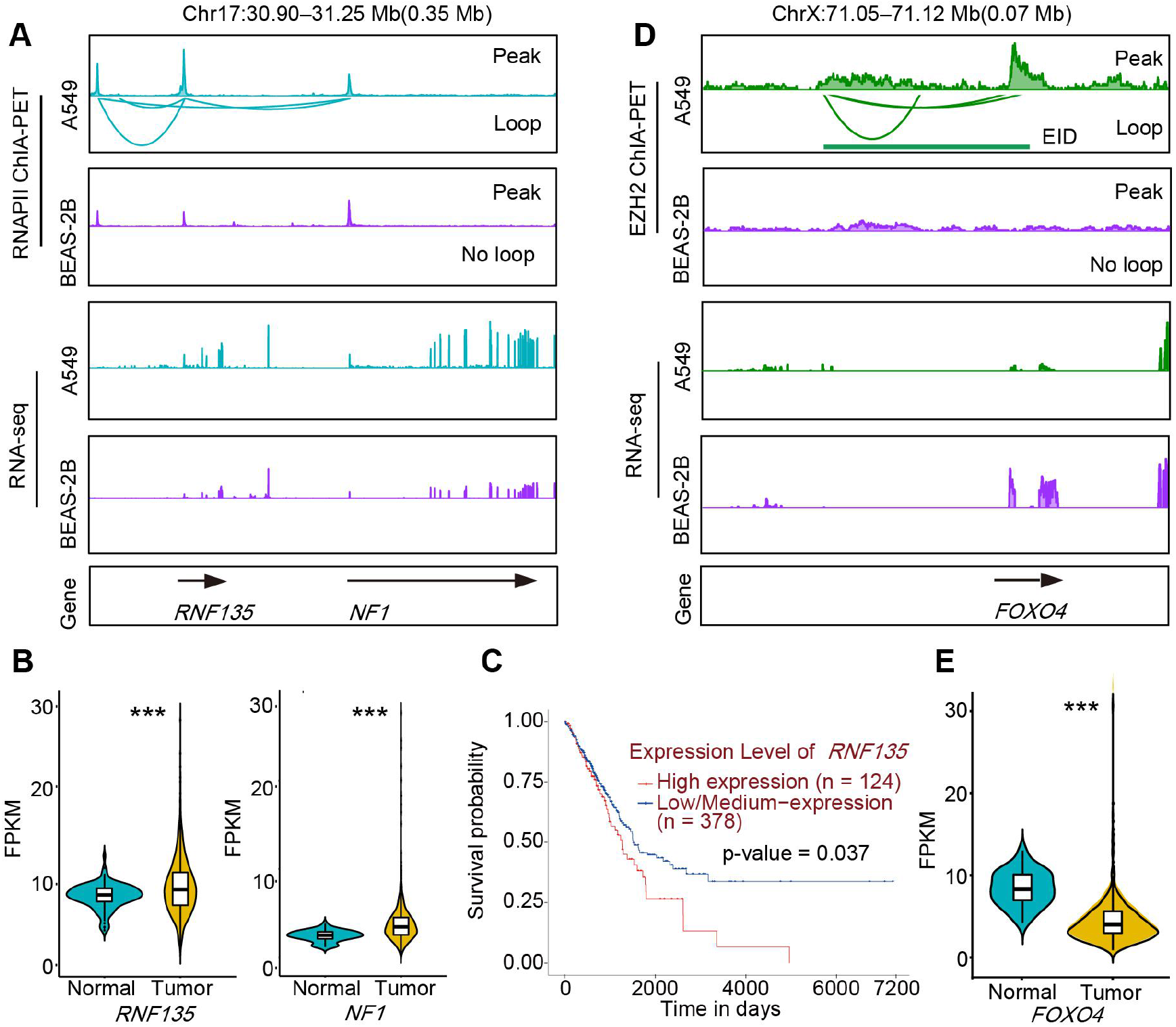
Differential interactions are associated with abnormal expression of oncogenes or tumor suppressor genes in lung cancer. **A.** *RNF135* and *NF1* promoter – promoter interactions that differ in lung cancer cell line A549 and noncancer cell line BEAS-2B. RNA-seq tracks show expression signals of genes from both cell lines. **B.** Violin plots overlaid with boxplots show distribution of *NF1* and *RNF135* mRNA expression levels in a large sample of LUAD tumor tissues and normal tissues from TCGA database. *P*-value was determined using one-sided Mann-Whitney U test. *** *P* < 0.001. **C.** Overall survival curve shows that overexpression of *RNF135* is correlated with poor outcome. This figure was made using online portal UALCAN [58]. Red survival curve is patients with *RNF135* gene expression in highest quartile, blue curve is for other samples with lower *RNF135* gene expression. **D.** EZH2-associated interactions on *FOXO4* gene promoter. EID marked in A549 loop track was identified by analysis. **E.** Violin plots overlaid with boxplots show distribution of *FOXO4* mRNA expression levels in a large set of LUAD tumor tissues and normal tissues from TCGA database. *P-*value was determined using one-sided Mann-Whitney U test. *** *P* < 0.001.

Abnormal repression of tumor suppressor genes is another crucial aspect of cancer [20]. In our study, several tumor suppressor genes were abnormally repressed by EZH2-associated loops. For example, *FBLN1* and *FOXO4*, well-known lung cancer suppressor genes [40–42], were significantly repressed (FPKM of *FBLN1* in A549: 13.02, BEAS-2B: 121.99; FPKM of *FOXO4* in A549: 0.53, BEAS-2B: 3.72) (Table S3) by EZH2-associated interactions in the A549 cells (Figure 5D, Figure S5A), and exhibited lower expression in LUAD samples than in normal samples (Figure 5E, Figure S5B). However, there were no EZH2-associated interactions in these two genes in the BEAS-2B cells. These results suggest that chromatin interactions are closely related to the expression levels of lung cancer-related genes.

To test whether cancer-specific interaction anchors can function as oncogenic regions, we performed CRISPR/Cas9-targeted knockout (KO) of the cancer-specific anchors to verify the effect of dysregulated interactions on cancer gene expression. *NCOA3* is a well-studied oncogene that is up-regulated in many cancers [43, 44] . In this study, the expression level of *NCOA3* was much higher in the A549 cells than in the BEAS-2B cells. In the A549 cells, there was a specific RNAPII mediated enhancer–promoter interaction cluster between the promoter of *NCOA3* and enhancer located in the first intron of *NCOA3* (**Figure 6A**), which did not exist in the BEAS-2B cells. We deleted (knocked out) the 1-kb enhancer region in the A549 cells (see Figure 6B) and characterized gene expression changes (Figure 6C). Results showed that expression was significantly lower in the KO A549 cells than in the wild-type A549 cells. *NCOA3* is known to have a positive effect on tumor cell growth [43, 44]. Thus, our cell growth study indicated that KO of cancer-specific interactions may reduce the cell growth rate (Figure 6D).

**Figure 6.**
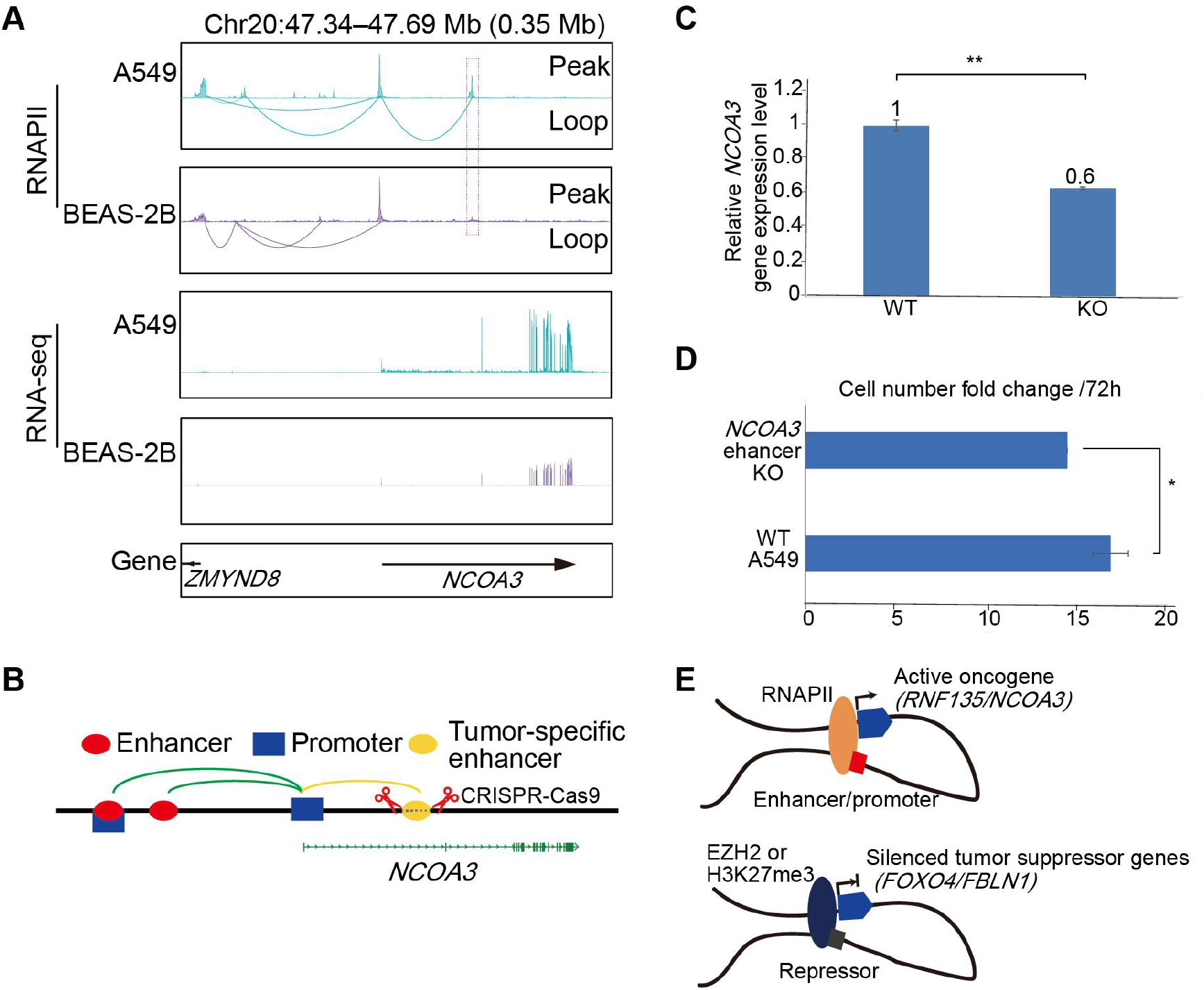
Knockout of lung cancer-specific anchor can reduce abnormal expression of oncogene *NCOA3*. **A.** Peak, loops, and RNA signal tracks around oncogene *NCOA3*, with a cancer-specific interaction linking intron enhancer and promoter of *NCOA3*. RNA-seq tracks show expression signals of genes from both cell lines. **B.** Schematic of KO of interaction anchor. KO of cancer-specific enhancer reduced long-range dysregulation of *NCOA3* promoter. **C.** *NCOA3* gene expression levels were detected by real-time qPCR. We used three biological RNA samples of *NCOA3* KO cells. *P*-value was determined using two-sided unpaired *t*-test. ** *P* < 0.01. **D.** Cell growth rate, indicated by number of cells, increased over 72 h. We calculated fold-change in cell growth in three sets of 6-well plates. KO of cancer-specific interactions reduced cell growth rate. *P*-value was determined using two-sided unpaired *t*-test. * *P* < 0.1. **E.** Models show function of active chromatin interactions in the regulation of oncogenes (upper) and function of abnormal repressive chromatin interactions in the regulation of tumor suppressor genes (low).

Based on the above results, we proposed two models as shown in Figure 6E, that describe the effects of abnormal interactions in lung cancer in terms of activating oncogenes and suppressing tumor suppressor genes. In lung cancer cells, some tumor suppressor genes were silenced by repressive factor-associated interactions. The differential expression patterns of these genes in cancer and noncancer tissues were non-significant in some types of cancer (e.g., breast cancer and esophageal carcinoma) (Figure S5C and D), indicating that dysregulation of these genes may be specific to LUAD. These dysregulated interaction patterns may be an important aspect of tumorigenesis and highlight the importance of normal interactions in cells.

## Discussion

In this study, we clarified the landscape of interactions associated with different factors in lung cancer cells. We found that the combination of interactomes associated with CTCF, RNAPII, and repressive factors can form a comprehensive 3D structure map of the genome, similar to that obtained with Hi-C. This similarity suggests that repressive interactions are an essential part of 3D genome structures.

The interactomes associated with various factors can be observed and analyzed separately. However, few studies on repressed genome structures in cancer have been reported. Here, we conducted innovative research on the interactions associated with repressive factors. We demonstrated that different types of interactions exhibit different distributions along the genome and different regulatory functions in the genome. We also identified several features of EZH2- and H3K27me3-associated interactions. First, repressed chromatin interactions tended to be weak, and most interactions failed to form clusters with high PET counts between interacting regions. Second, the EZH2- and H3K27me3-associated interaction anchors were widely distributed across large areas. Third, in most cases, repressive and active interactions were distributed in different genomic regions at intervals. Last, many genes were largely repressed by interactions with repressors. The repressive interactions primarily linked repressive elements to silenced or repressed genes. Loops may therefore be an additional pathway for the repression of genes.

Factor-enriched ChIA-PET can vastly increase the resolution and link regulatory elements to genes of interest [21, 22]. High-resolution loops can even distinguish between multiple TSSs of individual genes. Therefore, the ChIA-PET method has better practicality and deserves greater attention in cancer genome research. In the current study, the high-resolution ChIA-PET data revealed 10-bp and 190-bp signals, representing the DNA double helix turn and mononucleosome distance, respectively. Interestingly, the 3–4 nucleosome signals in the facultative heterochromatin suggests the structure mentioned in previous studies in the context of chromatin fiber packaging.

We used high-resolution chromatin interactions to map the whole-genome regulatory relationship and coexpression patterns of cancer-related genes and SNPs, which should expand current knowledge of regulatory networks in lung cancer. In future work, we will identify long-range regulatory elements and link mutation sites to target genes. As tumor-related genes and fusion transcripts were enriched in the interaction loops, we speculate the central role of long-range interactions in tumorigenesis.

Based on the comparison of cancer and noncancer interactomes, we identified many changed loops associated with the expression of cancer-related genes, including some that may play critical roles in lung cancer tumorigenesis. The abnormal formation of interaction clusters between the NF1 and RNF135 gene promoters increased the expression level of both genes, which has not been reported previously. Both expression and variance of these two genes was higher in TCGA LUAD samples than in normal samples. High RNF135 gene expression was specific to LUAD (Figure 5B, Figure S5C and D). Moreover, survival analysis suggested poor outcome in patients with *RNF135* overexpression. As shown in Figure S3B, loss of the EZH2-mediated interaction with *DKK1* may contribute to the up-regulation of this survival-related gene in A549 cells. Thus, these differential interactions may also be the cause or a symptom of dysregulation of cancer-related gene expression in cancer cells. Our results revealed a potential molecular marker and pathogenesis of cancer. The detection of abnormal interactions between cancer-related genes may be a novel diagnostic approach and powerful method for identifying oncogenes.

In conclusion, we mapped the high-resolution genome-wide interactomes mediated by CTCF, RNAPII, EZH2, and H3K27me3. These factors exhibited distinct distributions along the genome and distinct regulatory functions, and their combined interactomes largely represented the whole-genome interactome. By integrating RNA-seq and epigenomic data, we successfully described the relationship between gene expression, 3D genome interactions, interaction domains, and epigenetic modifications. We determined the interaction domain features and additional silencing function of repressive interactions and further described how abnormal interactions can dysregulate the expression of tumor-related genes. The transcription regulation and gene silencing networks generated from the ChIA-PET data will expand our understanding of the lung cancer genome and broaden the landscape of lung cancer research.

## Materials and Methods

### Cell culture

The A549 cell line (accession number CCL-185) was purchased from the American Type Tissue Culture Collection (ATCC) and was authenticated by the GENEWIZ Company. The BEAS-2B cell line was donated by Dr. Honglin Jin’s laboratory at the Cancer Center of Union Hospital, Tongji Medical College, Huazhong University of Science and Technology. The A549 and BEAS-2B cell lines were cultured at 37 °C under 5% CO_2_ in air. The A549 cells were cultured in Ham’s F12K medium (Cat. No. 21127022, Gibco, Grand Island, NY, USA) supplemented with 10% fetal bovine serum (FBS; Cat. No. 10099141, Gibco), 0.1 mM non-essential amino acids (Cat. No. 11140050, Gibco), and 100 U ml^-1^ penicillin/streptomycin (Cat. No. 15140163, Gibco). The BEAS-2B cells were cultured in RPMI 1640 (Cat. No. 11875101, Gibco) supplemented with 10% FBS and 100 U ml^-1^ penicillin/streptomycin. For both cell lines, medium was changed every day. The cells were passaged by trypsin digestion three times per week.

### Long-read ChIA-PET

For long-range interaction analysis of the A549 and BEAS-2B cells, we used long-read ChIA-PET as described previously [21] with minor modifications. The cells were resuspended using trypsin digestion and then dual cross-linked with 1.5 mM ethylene glycol bis (succinimidyl succinate) (Thermo Fisher Scientific, 21565) for 40 min and with 1% formaldehyde (Sigma, F8775) for 10 min at room temperature. The cells were lysed, and chromatin was fragmented into 1–5-kb fragments by sonication (high level, 33 cycles, 30 s ON, 50 s OFF) using a Bioruptor (Diagenode). Chromatin immunoprecipitation was used to enrich the complex fragments with magnetic beads of protein G (Thermo) and 60–100 µg of antibodies against RNAPII (Santa SC-56767), EZH2 (CST 5246), H3K27me3 (ABclonal custom-made), and CTCF (ABclonal A1133). The beads were then washed, and DNA blunt ends were prepared with A tails. Bridge linkers were used to ligate the proximal DNA ends in a 1.8-ml reaction system using T4 DNA ligase (Invitrogen). The following DNA extraction and library construction steps were the same as the previous protocol. The libraries were paired-end sequenced (2 × 150 bp) using the Illumina HiSeq X Ten system.

### Total RNA-seq

For RNA-seq, total RNA was extracted from 1 million cells using RNeasy columns (QIAGEN). Ribosomal RNA (rRNA) was depleted using an rRNA Depletion Kit (NEB), and then strand-specific libraries were constructed using the NEBNext Ultra II RNA Library Prep Kit for Illumina. Paired-end (2 × 150 bp) sequencing of libraries was conducted using the Illumina HiSeq X Ten system.

### CRISPR/Cas9 KO of enhancer

For the KO system, we applied chromatin fragment deletion as described previously with some modifications [45]. The single-guide RNA (sgRNA) templates were constructed and incorporated into pGL3-U6-sgRNA-PGK-Puro plasmids. The A549 cells were cultured to approximately 60% confluence and transfected with Lipofectamine 3000 (Invitrogen) in a 6-well plate with 2 μg of pcDNA3.1-Cas9 and 1.5 μg of sgRNA plasmids for the upstream target and 1.5 μg for the downstream target. Puromycin was added two days later to a final concentration of 1.2 μg/ml. The cells were then diluted and plated in 15-cm plates and cultured for 5 days to isolate single clones. For KO of the NCOA3-specific enhancer, we designed guide RNA sequence: upstream 1: ATAGAATTGCAACCTCATGGAGG, upstream 2: CAGTTGTACCTACTGCCAAATGG, downstream 1: CCCCCTTTGGCTGAGATAAATGG, downstream 2: GTCTAGCACAATGTGGCACATGG.

### RNA expression analysis by quantitative real-time polymerase chain reaction (qRT-PCR)

QRT-PCR was conducted on a CFX-connect RT PCR detection system (Bio-Rad). The reaction used Genious 2X SYBR Green Fast qPCR Mix (ABclonal) for qRT-PCR. Total RNA was extracted from 1 million cells by RNeasy columns (QIAGEN). We used 2 µg of RNA to perform reverse transcription with random primers using TransScript One-Step gDNA Removal and cDNA Synthesis SuperMix TransGen Biotech. The qRT-PCR was then performed using primers for *NCOA3* mRNA (primers of *NCOA3* qRT-PCR forward: GGACCTGGTTAACACAAGTG and reverse: GTCCAGGAAACTCCATTAACTG).

### Analysis of ChIA-PET data

Long-read ChIA-PET sequence data were obtained using a modified ChIA-PET Tool pipeline [46]. Briefly, after trimming the linkers, the sequences flanking the linker were mapped to the human reference genome (hg38) using bwa-mem [47], and only uniquely mapped (MAPQ >= 30) PETs were retained. Each PET was categorized as either a self-ligation PET (two ends of the same DNA fragment, with a genomic span less than 8 kb) or an inter-ligation PET (two ends from two different DNA fragments in the same chromatin complex from different chromosomes, or from the same chromosome with a genomic span of more than 8 kb). Self-ligation PETs were used for binding site calling and inter-ligation PETs were used for long-range interaction calling. For RNAPII and CTCF, we used centered 3-kb regions of RNAPII and CTCF chromatin immunoprecipitation with sequencing (ChIP-seq) peaks (GSE31477 [48]) as given anchors to call interaction clusters. To obtain high-confidence interactions, we included the detected interactions only if the false discovery rate (FDR) was less than 0.05 and the PET count was three or more. To evaluate the robustness of the ChIA-PET method, we analyzed the biological replicates of the ChIA-PET libraries at several different resolutions. Some analysis results are presented in Figure S1. Moreover, we transformed unique ChIA-PET mapping reads to a contact matrix using ChIA-PET2 (version 0.9.3) [49] software and normalized the matrix using the iterative correction method from HiC-Pro (version 2.11.1) [50] software. We also used Juicer (version 1.7.6) [51] software to calculate A/B compartments at 1-Mb and 100-kb resolutions. When calculating Pearson correlation coefficients of the four factor-mediated contact and Hi-C matrices, we used Hi-C data from the ENCODE database (accession number ENCSR662QKG).

### Hierarchical chromatin structure analysis

We calculated the genomic spans of the self-ligation and inter-ligation PETs and observed 10-bp and 190-bp signals, respectively. We speculated that the two signals were DNA double helices and nucleosome units, respectively. To observe the tetranucleosome signal, we defined CTRs (center 2 kb of overlapping regions of H3K9me3 and H3K4me3 peaks) between euchromatin and heterochromatin, H3K9me3 and H3K4me3 ChIP-seq data were obtained from the GEO database (accession numbers GSE29611). Nearly 11 003 CTRs were obtained. We choose inter-ligation PETs in CTRs and calculated their genomic span.

### Annotation of interaction loops

We annotated loops using gene promoter, enhancer, and repressor information. The promoters were defined as the ±2-kb regions among the TSSs. Cell-type-specific enhancer and repressor annotations were adopted from the A549 ChromHMM data [52], and both strong and weak/poised enhancers/repressors were used. We classified loops according to the overlap of interaction anchors with promoters, enhancers, or repressors, with priority given to the promoter region. For example, we defined promoter – promoter loops as both interaction anchors overlapping with promoters, promoter– enhancer loops as one anchor overlapping with a promoter and the other anchor overlapping with an enhancer, and promoter– repressor loops as one anchor overlapping with a promoter and the other anchor overlapping with a repressor. We further categorized the promoter–enhancer and promoter–repressor loops according to enhancer/repressor locations in relation to the gene body: i.e., intragenic proximal enhancers/repressors, which are located inside a gene body and interact with the nearest promoters; extragenic proximal repressors/enhancers, which are located outside a gene body and interact with the nearest promoters; intragenic distal enhancers/repressors, which are located inside a gene body, bypass nearby genes, and interact with gene promoters over long distances; and extragenic distal enhancers/repressors, which are located outside all gene bodies, bypass nearby genes, and interact with gene promoters over long distances.

### Coexpression analysis of genes with promoter – promoter interactions by Pearson correlation coefficients

To investigate the coexpression of genes with CTCF, RNAPII, EZH2, and H3K27me3 promoter – promoter interactions defined by ChIA-PET, we calculated the Pearson correlation coefficients of expression levels of gene pairs with promoter – promoter interactions. Gene expression data for 535 LUAD samples were downloaded from the TCGA. The genes involved in promoter – promoter interactions were randomly rewired to compile a random control with the same gene background but different pairings. Randomly selected gene pairs with similar distributions of genomic span and gene density as the promoter–promoter regions were used as additional controls.

### Analysis of chromatin interaction domains

A chromatin interaction domain is a genomic region spanned by continuously connected loops mediated by specific factors. First, based on the continuous connectivity of loops, we clustered loops into candidate domains. Second, we calculated loop coverage along all chromosomes at base-pair resolution and subtracted low-coverage regions from the candidate domains. Finally, the interaction domains with genomic size smaller than 10 kb were excluded, and the final EIDs, HIDs, and RIDs were obtained. When comparing histone modifications of different domains, we used ChIP-seq data (H3K27ac, H3K4me3, H3K27me3, and H3K9me3) obtained from the GEO database (accession numbers GSE29611 and GSE32465).

### RNA-seq data processing

Strand-specific poly(A) RNA-seq libraries were generated and sequenced as 150-bp paired-end reads. For RNA-seq analysis, we conducted quality control using FastQC (https://www.bioinformatics.babraham.ac.uk/projects/fastqc/), and adaptor sequences were removed using Trimmomatic (version 0.32) [53]. After quality filtering, the reads were mapped to the human reference genome (hg38) by hisat2 (version 2.1.0) [54], and gene expression was qualified by StringTie (version 1.3.4d) [55]. Differential gene expression was calculated using the DESeq2 [56] package in R. To identify significant differentially expressed genes between the A549 and BEAS-2B cell lines, we used thresholds of adjusted *P*-value < 0.01 and absolute log2-fold-change > 1 in expression levels.

### Gene expression at loop anchors

We categorized genes into four types according to whether their gene promoters were associated with RNAPII loops or H3K27me3 loops: 1) genes whose promoters were only associated with RNAPII loops but not associated with H3K27me3 loops; 2) genes whose promoters were associated with both RNAPII loops and H3K27me3 loops; 3) genes whose promoters were neither associated with RNAPII loops nor H3K27me3 loops; and 4) genes whose promoters were not associated with RNAPII loops but were associated with H3K27me3 loops. Significant differences between the FPKM values of these different categories of genes were compared using Wilcoxon rank-sum tests. Similarly, we categorized genes into four types according to whether their promoters were associated with EZH2/RNAPII loops or EZH2/RNAPII binding sites and compared expression differences between groups.

### Gene Ontology (GO) analysis

For genes in the cell-specific anchors with significantly differential expression, we examined enrichment of GO terms using the Database for Annotation, Visualization, and Integrated Discovery (DAVID) [57].

### TCGA RNA-seq data analysis

We downloaded gene expression data for 535 primary LUAD tumor tissues and 59 lung solid normal tissues from the TCGA database and analyzed their gene expression differences, especially for GWAS SNP target genes and lung cancer-related genes (*NF1*, *RNF135*, *FBLN1*, and *FOXO4*).

## Data availability

Sequence data generated in this study were deposited in the Genome Sequence Archive [52] in the National Genomics Data Center, Beijing Institute of Genomics, Chinese Academy of Sciences/China National Center for Bioinformation (GSA: HRA000295 with BioProject: PRJCA003299), and are publicly accessible at http://bigd.big.ac.cn/gsa-human and https://ngdc.cncb.ac.cn/bioproject/.

## Acknowledgments

This study was supported by a General Program of the National Natural Science Foundation of China (NSFC) (Grant No. 31970590). We thank Dr. Honglin Jin from Union Hospital, Tongji Medical College, Huazhong University of Science and Technology for the BEAS-2B cell line. We acknowledge the TCGA and ENCODE databases for providing their platforms and contributors for their meaningful datasets.

