## Supplementary File for "Mapping Multiple Factors Mediated Chromatin Interactions to Assess Regulatory Network and Dysregulation of Lung Cancer-Related Genes"

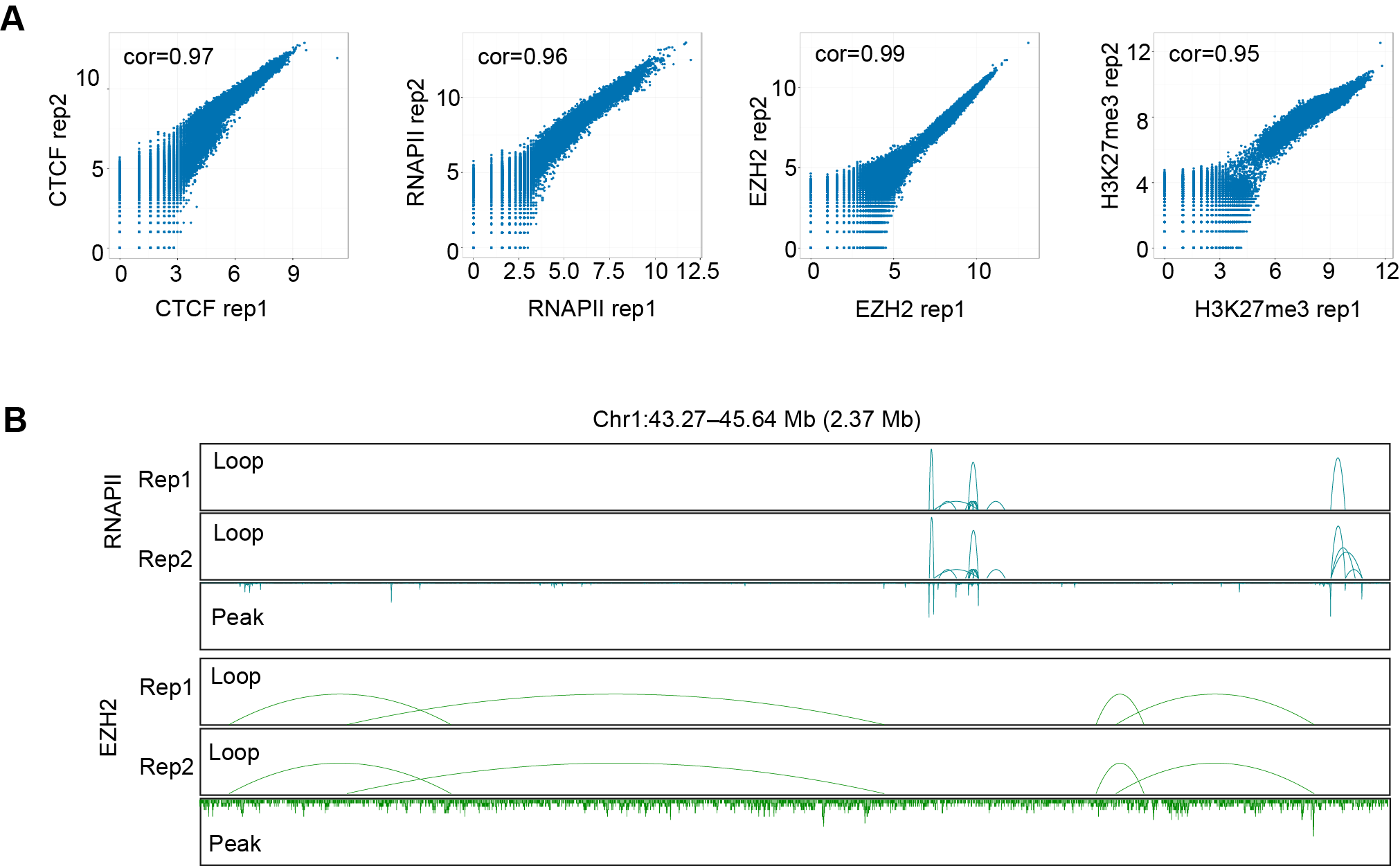

**Figure S1 Reproducibility of ChIA-PET data**

A. Scatter plots showing the contact matrix correlation between different ChIA-PET replicates in A549 cells. **B.** Comparison of ChIA-PET interaction clusters between two RNAPII replicates and two EZH2 replicates represented by chromosome 1: 43.27–45.64 Mb.

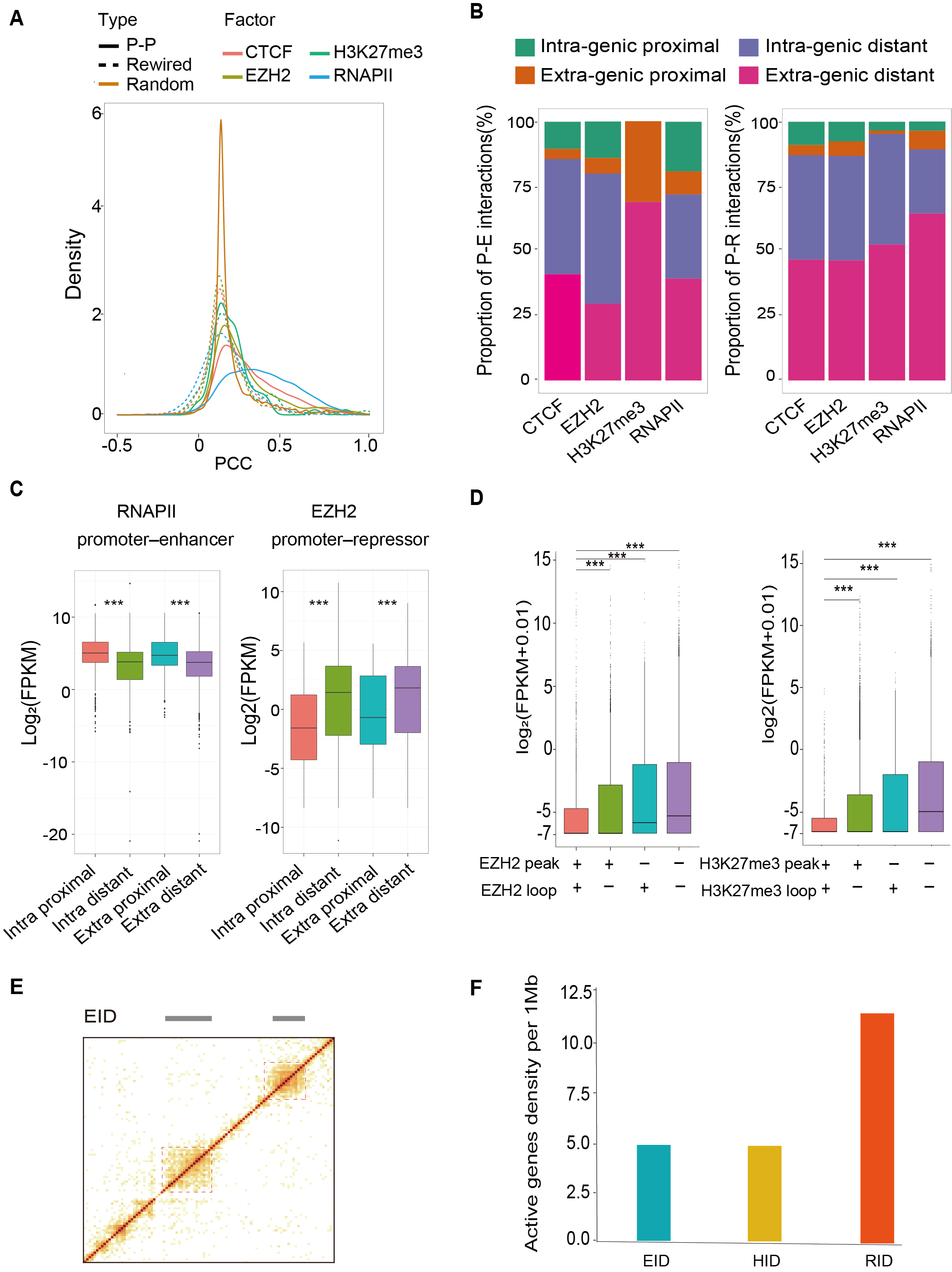

**Figure S2 Factor-specific interaction domains are important gene-regulatory units in the genome**

A. Distribution of Pearson correlation coefficient values for gene pairs with promoter–promoter interactions, randomly rewired gene pairs, and randomly picked gene pairs from control regions with the same genomic span. P–P represents gene pairs with promoter-promoter interactions. **B.** Proportional distribution of 4 classes of enhancers and repressors observed in A549 cells based on locations concerning gene coding regions. "Intra-genic proximal" enhancers are located inside the gene body and interact with nearby promoters. "Extra-genic proximal" enhancers are located outside of the gene body and interact with nearby promoters. "Intra-genic distal" enhancers are located inside the gene body, bypass nearby genes and interact with faraway gene promoters over long distances. "Extra-genic distal" enhancers are located outside of the gene body, bypass nearby genes and interact with faraway gene promoters over long distances. **C.** Comparison of the expression levels of genes with different types of promoter–enhancer or promoter–repressor interactions. The p-value was determined using one-sided Mann-Whitney U test. *** for *P* < 0.001. **D.** Expression levels of genes whose promoters are with or without EZH2 or H3K27me3 binding or loop anchor binding. Mann-Whitney U test was used to calculate difference significance, *** for *P* < 0.001. "+" means that the genes are located in the peak regions or loop anchor regions, and "-" means they are not. **E.** Examples of EZH2 interaction domains are showed in the heatmap. **F.** Densities of active genes (FPKM > 1) per 1 Mb in the EIDs, HIDs, and RIDs.

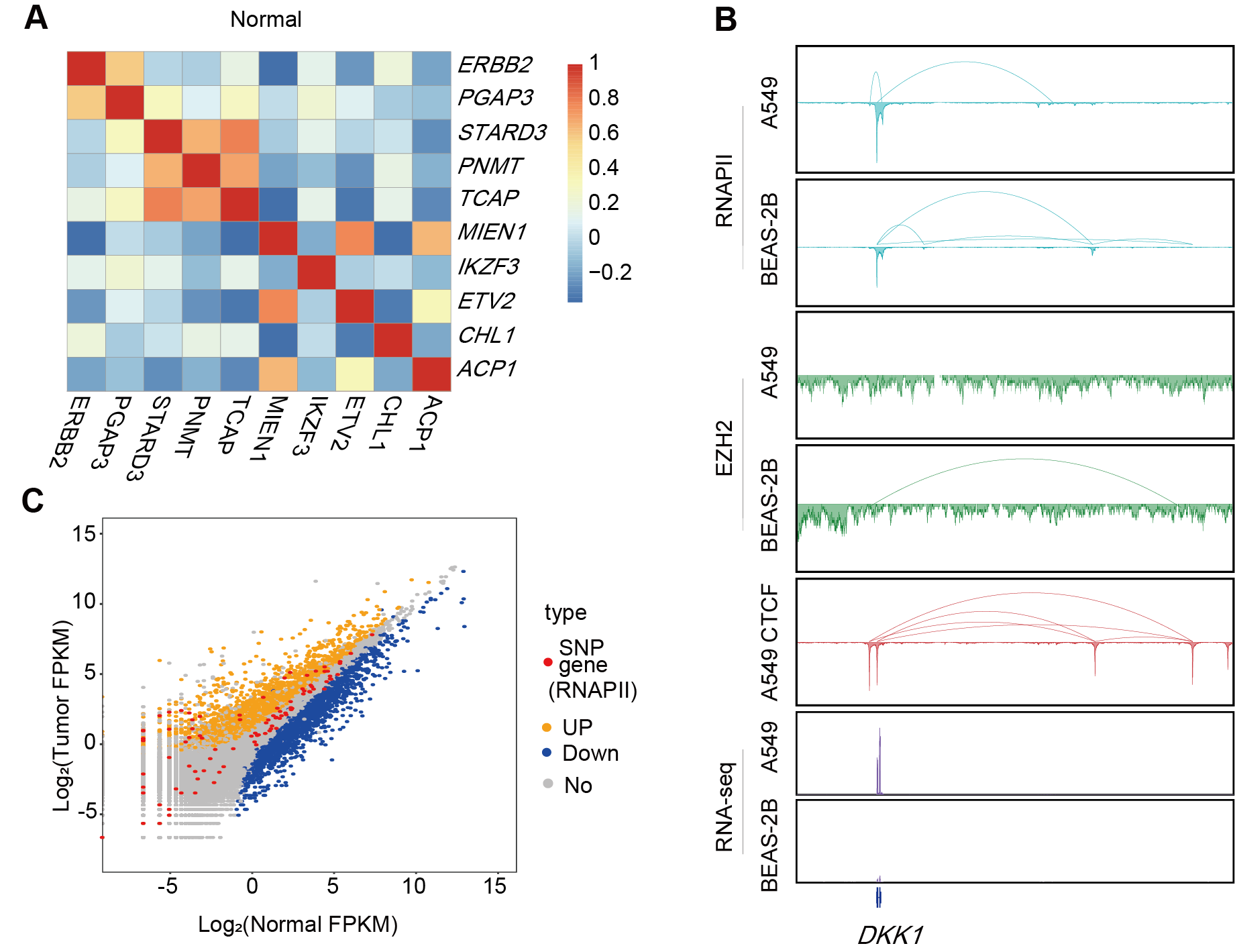

**Figure S3 The regulatory network of lung cancer-related genes and SNPs**

A. The Pearson correlation coefficients for *ERBB2* versus interacting genes in normal samples from the TCGA LUAD dataset. The genes are the same as those in Figure 4C. The coexpression pattern was much weaker in the normal samples. **B.** The interaction loops on the survival-related gene *DKK1* in A549 and BEAS-2B cells. The upper track in each box shows the interaction clusters associated with each factor. The peak density in the lower track shows the ChIP-seq signals in the cells. The RNA-seq signals are shown in the bottom tracks. The additional interaction associated with EZH2 in BEAS-2B cells may contribute to the decreased expression of *DKK1* in normal cells. **C.** Comparison of gene expression in tumor and normal samples from the TCGA LUAD dataset. The genes interacting with lung cancer risk-related SNPs are marked as red dots. Genes that are not differentially expressed are marked as gray dots. Upregulated genes in the tumor samples are marked as yellow dots, and downregulated genes in the tumor samples are marked as blue dots.

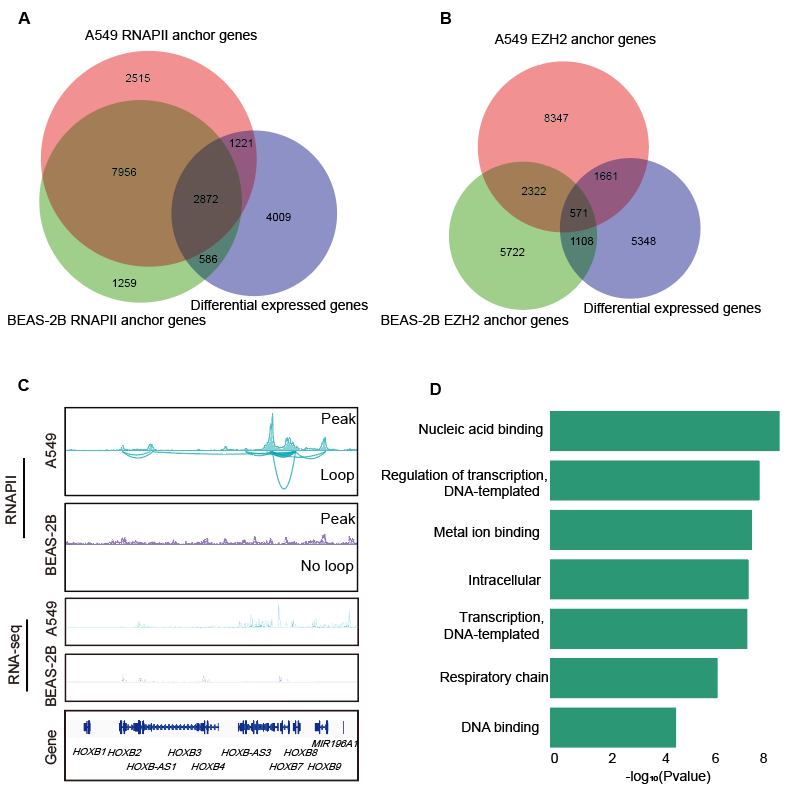
**Figure S4 Genes with different interactions and expression in A549 and BEAS-2B cell lines**

A-**B**. Venn diagrams of genes with chromatin interactions located in their promoter regions. **A.** Genes with interactions associated with RNAPII. **B**. Genes with interactions associated with EZH2. The red circles represent the genes with specified chromatin interactions in A549 cells. The green circles represent the genes with specified chromatin interactions in BEAS-2B cells. The purple circles represent the genes with significantly different expression levels (p-value < 0.01, fold change > 2) in the two cell lines. **C.** An interaction domain around the *HOXB* gene cluster on chromosome 17 exists specifically in A549 cells. The lower track in each box shows the interaction clusters associated with RNAPII among *HOXB* genes. The peak density in the upper track shows the RNAPII binding profile in the two cell lines. The RNA-seq tracks show the RNA signals. **D.** GO analysis terms of specific RNAPII interactions and differentially expressed genes in BEAS-2B cells.

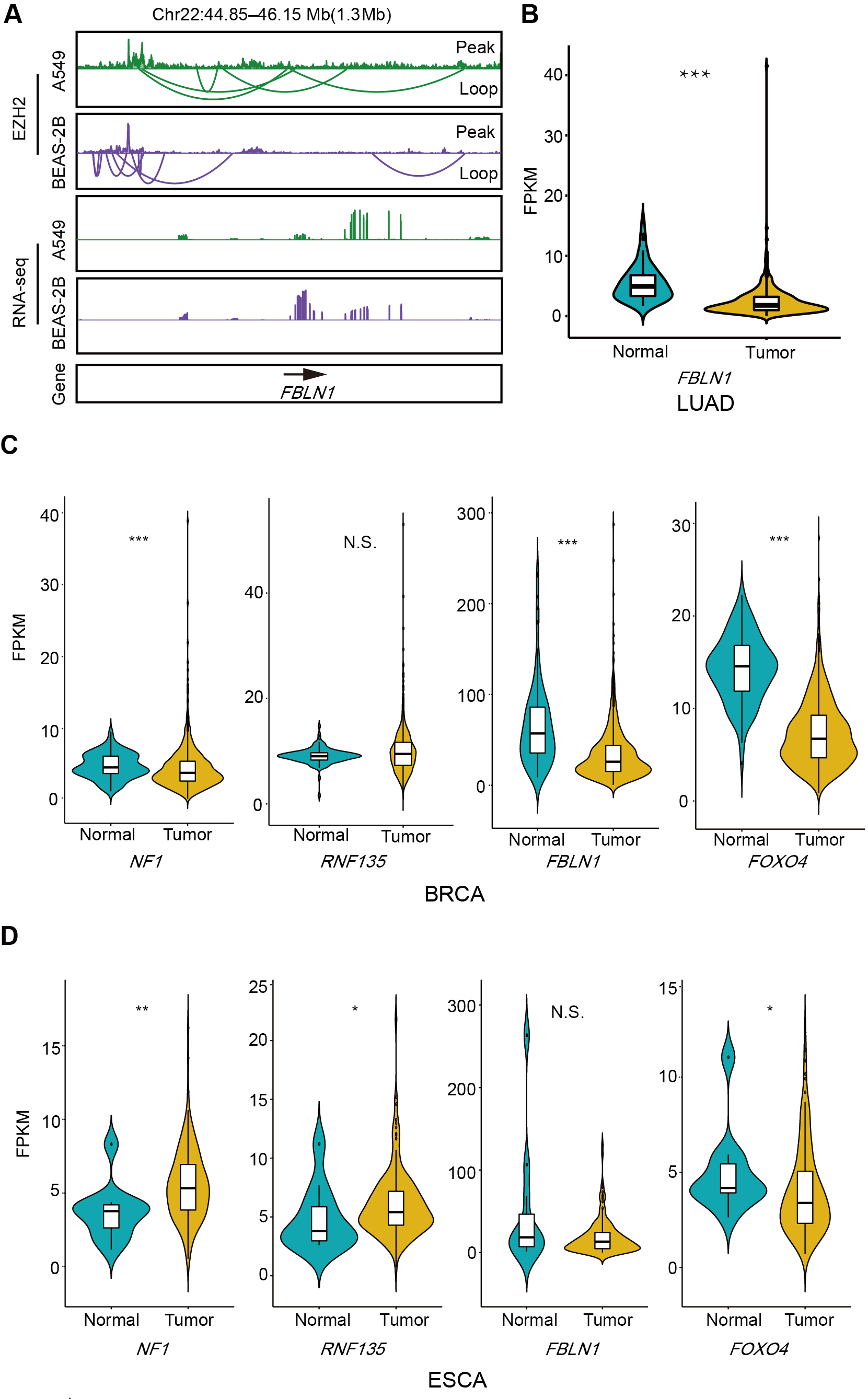

**Figure S5 Expression profiles of four cancer-related genes in the TCGA database**

**A.** Example of the EZH2-associated interactions on the FBLN1 gene promoter that are different in the lung cancer cell line A549 and noncancer cell line BEAS-2B. The RNA-seq tracks show the expression signal of FBLN1 in the two cell lines. **B.** Violin plots overlaid with boxplots show the distribution of FBLN1 mRNA	expression levels in a large set of LUAD tumor tissues and normal tissues from the TCGA database. The p-value was determined using a one-sided Mann–Whitney U test. *** for *P* < 0.001. **C.** Violin plots overlaid with boxplots show the distributions of NF1, RNF135, FBLN1, and FOXO4 mRNA expression levels in a large group of breast cancer (BRCA) tumor tissues and normal tissues from the TCGA database. The p-value was determined using a one-sided Mann-Whitney U test. *** for *P* < 0.001; ** for *P* < 0.01; N.S. means no significant difference. The upregulation of RNF135 in LUAD was not significant in BRCA samples. The NF1 has opposite expression patterns in breast cancer. **D.** Violin plots overlaid with boxplots show the distributions of NF1, RNF135, FBLN1, and FOXO4 mRNA expression levels in a large set of esophageal carcinoma (ESCA) tumor tissues and normal tissues from the TCGA database. The p-value was determined using a one-sided Mann-Whitney U test. ** for *P* < 0.01; * for *P* < 0.1; N.S. means no significant difference.
